# Neural computation underlying rapid learning and dynamic memory recall for sensori-motor control in insects

**DOI:** 10.1101/2020.04.05.026203

**Authors:** Hannes Rapp, Martin Paul Nawrot

## Abstract

Foraging is a vital behavioral task for living organisms. Behavioral strategies and abstract mathematical models thereof have been described in detail for various species. To explore the link between underlying nervous systems and abstract computational principles we present how a biologically detailed neural circuit model of the insect mushroom body implements sensory processing, learning and motor control. We focus on cast & surge strategies employed by flying insects when foraging within turbulent odor plumes. Using a synaptic plasticity rule the model rapidly learns to associate individual olfactory sensory cues paired with food in a classical conditioning paradigm. Without retraining, the system dynamically recalls memories to detect relevant cues in complex sensory scenes. Accumulation of this sensory evidence on short timescales generates cast & surge motor commands. Our systems approach is generic and predicts that population sparseness facilitates learning, while temporal sparseness is required for dynamic memory recall and precise behavioral control.

## 1. Introduction

Navigating towards a food source during foraging requires dynamical sensory processing, accumulation of sensory evidence and appropriate high level motor control. Navigation based on an animals’ olfactory sense is a challenging task due to the complex spatiotemporal landscape of odor molecules. A core aspect of foraging is the acquisition of sensory cue samples in the natural environment where odor concentrations vary rapidly and steeply across space. Experimental access to the neural substrate is challenging in freely behaving insects. Biologically realistic models thus play a key role in investigating the relevant computational mechanisms. Consequently, recent efforts at understanding foraging behavior have focused on identifying viable computational strategies for making navigation decisions (Vergassola et al., 2007; Masson, 2013; Gaudry et al., 2012; Hein et al., 2016; Baddeley et al., 2009; Baker et al., 2018; Cardé and Willis, 2008).

An odor plume is often considered a volume wherein odor concentration is generally above some behavioral threshold. At macroscopic scales and in a natural environment, however, plumes are turbulent (Murlis et al., 1992; Crimaldi et al., 2002). In turbulent conditions a plume breaks up into complex and intermittent filamentous structures that are interspersed with clean air pockets or below behavioral threshold concentration patches (Celani, 2014; Kree et al., 2013; Connor et al., 2018). The dispersing filaments form the cone-like shape of the macroscopic plume where the origin of the cone yields the position of the odor source. When entering the cone, flying insects encounter odor filaments as discrete, short-lived sensory events in time.

Several features have been derived from the statistics of an odor plume that provide information regarding the location of the odor source (Balkovsky and Shraiman, 2002; Celani, 2014; Crimaldi et al., 2002; Murlis et al., 2000; Shraiman and Siggia, 2000). The mean concentration varies smoothly in lateral and longitudinal directions of time-averaged (and laminar) plumes. However, for behavioral strategies animals cannot afford the time it takes to obtain stable macroscopic estimates of mean concentrations (Murlis et al., 1992). Vickers et al. (2001) and Park et al. (2016) proposed the time interval between odor encounters as an informative olfactory feature while Crimaldi et al. (2002) suggested intermittency, the probability of the odor concentration being above some behavioral threshold, as the relevant feature. However, similarly to estimating mean concentration, acquiring a sufficient number of samples for stable estimates of these quantities exceeds the time typically used to form behavioral decisions (Murlis et al., 1992). Hence, obtaining time averaged quantities is not an optimal strategy to guide navigation decisions as concluded by Boie et al. (2018).

Most animals perform searches at large distances from the odor source where the intermittency of plumes becomes a more severe problem as available sensory cues become more sparse in space and time. Thus, strategies that exploit brief, localized sensory cues for navigation have been studied by several groups. One strategy for medium and long-range navigation that has consistently been observed across species of flying insects emerges from a sequence of chained sensorimotor reflexes: casting & surging (van Breugel et al., 2015; Gaudry et al., 2012). Encountering a whiff of odor triggers an upwind surge behavior, during which the insect travels parallel to the wind direction. After losing track of the plume it evokes a crosswind cast behavior, in which a flight path perpendicular to the direction of air flow is executed. Performing repeated casts by U-turning allows the insect to reenter and locate the plume in order to trigger the next upwind surge (Cardé and Willis, 2008; van Breugel and Dickinson, 2014; van Breugel et al., 2015; Vickers and Baker, 1994; Riffell et al., 2014; Pang et al., 2018; Budick and Dickinson, 2006). As the subject approaches the source it increasingly makes use of visual cues for navigation as the plume narrows down. (van Breugel and Dickinson, 2014; Saxena et al., 2018).

A number of studies have proposed abstract mathematical models for optimal search algorithms that assumed different types of relevant navigational cues. The infotaxis method proposed in Vergassola et al. (2007) depends on extensive memory and priors regarding a plume’s structure. Contrary, in van Breugel and Dickinson (2014) only local cues are used. A standard algorithm for navigational problems in robotics is simultaneous localisation and mapping (SLAM), which has been used in Baddeley et al. (2009) to study olfactory navigation in bumblebees. An algorithm that works without space perception has been proposed by Masson (2013) using a standardized projection of the probability of source position and minimization of a free energy along the trajectory. Finally, the work of Pang et al. (2018) compares several models and shows that it is difficult to discriminate between different models based on behavioral responses. A recent work by Boie et al. (2018) using information-theoretic analysis shows that plumes contain both, spatial and temporal information about the source’s position.

While all of these previous mathematical methods for olfactory search algorithms have proven to successfully solve this task based on the respective assumptions, they share the same major drawback: none of them uses the computational substrate of the brain, spiking neurons and networks thereof. Instead, all methods make heavy use of symbolic math and advanced mathematical concepts that are not available to the biological brain. It is further unclear how and to what extend these methods could be implemented or learned by the nervous system. Additionally, the problem of navigation and foraging is often considered as an isolated task, independent from sensory processing.

Our approach distills recent experimental results to formulate a biologically plausible and detailed spiking neural network model supporting adaptive foraging behavior. We thereby take advantage of the rapidly accumulating knowledge regarding the anatomy (e.g. (Aso et al., 2014a; Caron et al., 2013; Xu et al., 2020)) and neurophysiology (e.g. (Ito et al., 2008; Kazama and Wilson, 2009; Demmer and Kloppenburg, 2009; Szyszka et al., 2014; Inada et al., 2017; Egea-Weiss et al., 2018)) of insect olfaction and basic computational features (Litwin-Kumar et al., 2017; Kloppenburg and Nawrot, 2014; Betkiewicz et al., 2020). We follow the idea of compositionality, a widely used concept in mathematics, semantics and linguistics. According to this principle, the meaning of a complex expression is a function of the meanings of its constituent expressions (Frege principle (Hintikka, 1984)). In the present case of foraging and navigation this means dynamically recombining memories of individual (temporal and spatial) sensory cues present within a plume.

## 2. Results

We approach the problem of foraging by decomposition into four components: First, sensory processing with temporal sparse and population sparse coding in the mushroom body (MB). Second, associative learning for assigning a valence to individual odor identities. Third, the time-dependent detection of valenced cues resulting from encounters of discrete odor filaments to provide an ongoing and robust estimate of sensory cue evidence. Fourth, the translation into online motor command signals to drive appropriate behavior.

For sensory processing we use a three-layer spiking neural network model of the insect olfactory pathway (see Fig 1). The generic blueprint of the insect olfactory system is homologous across species and comprises three successive processing stages (see Methods for details): The periphery with olfactory receptor neurons (ORNs), the antennal lobe (AL) and the MB. Excitatory feed-forward connections across layers from ORNs to projection neurons (PNs), from ORNs to local interneuron (LNs), and from PNs to the MB Kenyon cells (KCs) are fixed. Lateral inhibition within the AL uses fixed synaptic weights from LNs to PNs. For neuron numbers and their connectivity patterns we here rely on the adult *Drosophila melanogaster* where anatomical knowledge is most complete (Turner et al., 2008; Takemura et al., 2017; Xu et al., 2020; Aso et al., 2014a). A single MB output neuron (MBON) receives input from all Kenyon cells and plasticity at the synapses between KCs and the MBON enable associative learning (Gütig, 2016; Rapp et al., 2020).

**Figure 1:**
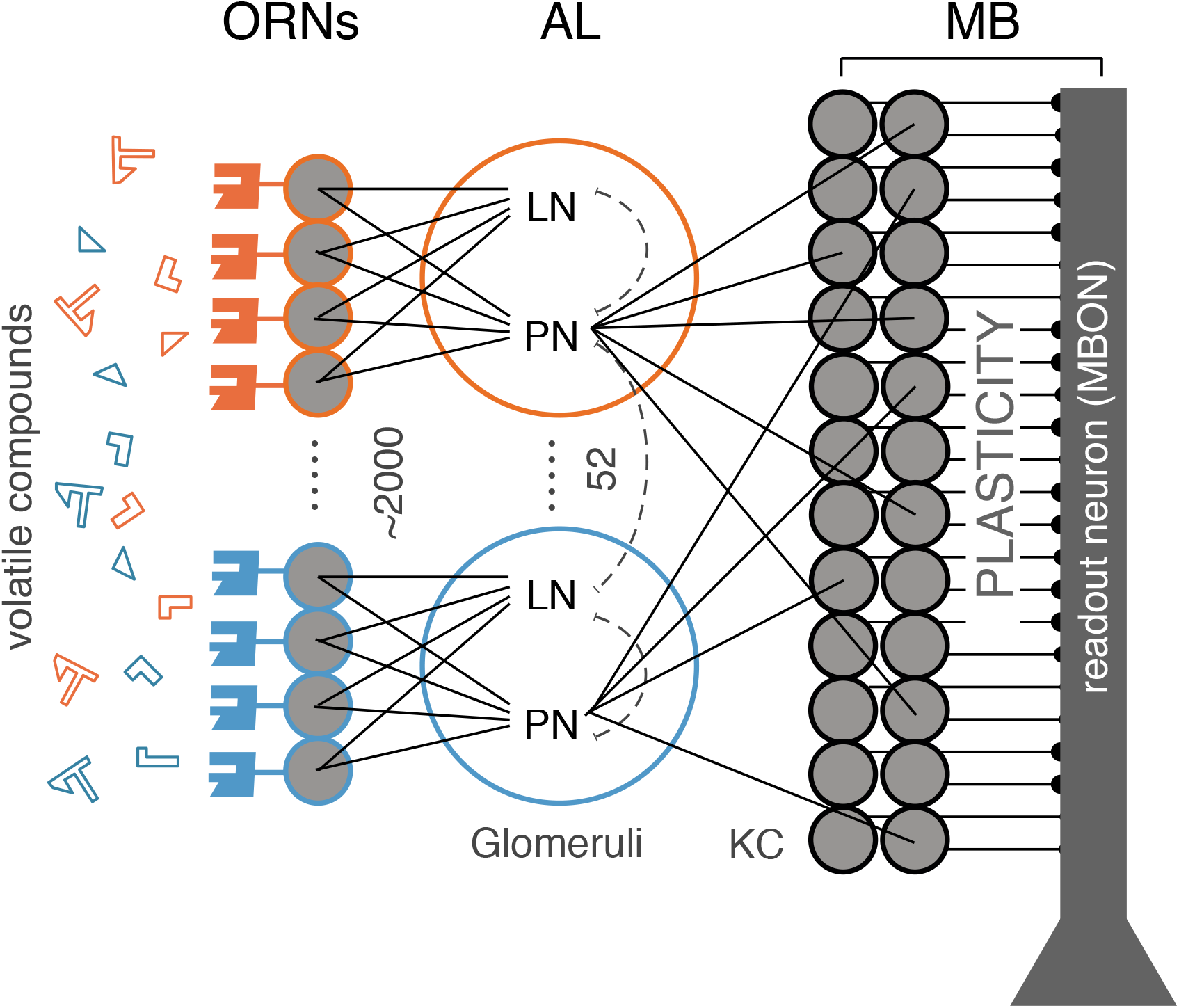
Spiking network model of the insect olfactory system. 2048 olfactory receptor neurons (ORNs) at the antennae bind and respond to volatile odorant compounds. ORNs expressing the same genetic phenotype (52 different receptor types) project to the same Glumerus in the antenal lobe (AL). Each of the 52 Glomeruli consitutes of one projection (PN) and local interneuron (LN). LNs form lateral inhibitory connections among Glomeruli and PNs randomly connect to a large population of Kenyon Cells (KC) where each KC receives input from 6 random PNs on average. Sensory processing, learning and memory is performed by the Mushroom body (MB) with Kenyon Cells (KC) and a fully connected single, plastic readout neuron (mushroom body output neuron (MBON)). The overall architectural bauplan of the olfactory system is homologous across species. Here the specific numbers of neurons within each population and connectivity are taken from the connectome of the mushroom body of the adult *Drosophila melanogaster*

### Sparse coding in space and time

The olfactory system transforms a dense olfactory code in the AL into a sparse stimulus code at the MB level. In the large population of KCs, a specific odor stimulus is represented by only a small fraction of all KCs (population sparseness) and each stimulus-activated KC responds with only a single or very few action potentials (temporal sparseness).

In our model, temporal sparseness is achieved through the cellular mechanisms of spike-frequency adaptation (SFA, Benda and Herz (2003); Farkhooi et al. (2013); Betkiewicz et al. (2020)) implemented at two levels of the system. ORNs show clear stimulus response adaptation that could be attributed to the spike generating mechanism (Nagel and Wilson, 2011). Based on this experimental evidence we introduced a slow and weak SFA conductance in our model ORNs (see Methods). At the level of the MB, KCs have been shown to express strong SFA-mediating channels (Demmer and Kloppenburg, 2009). This is matched by the SFA parameters of our model KCs (see Methods, (Farkhooi et al., 2013; Betkiewicz et al., 2020)). As an effect of cellular adaptation in ORNs and KCs, odor stimulation (Fig 2 A) results in temporally precise and adaptive responses across all layers of the network (Fig 2B). The effect of SFA implemented in ORNs is transitive and thus carries over to the postsynaptic PN and LN populations in agreement with experimental observations across species (Stopfer et al., 2003; Wilson et al., 2004; Bhandawat et al., 2007; Krofczik et al., 2009; Watanabe et al., 2012). In the KC population the background firing rate is very low. This is partially due to the outward SFA conductance (Farkhooi et al., 2013) and in agreement with experimental results (Ito et al., 2008). The KC population response is highly transitive where individual responding cells generate only a single or very few response spikes shortly after stimulus onset. This is in good qualitative and quantitative agreement with the temporal sparse KC responses measured in various species (Perez-Orive et al., 2002; Stopfer et al., 2003; Ito et al., 2008; Turner et al., 2008; Gruntman and Turner, 2013; Szyszka et al., 2005; Froese et al., 2014).

**Figure 2:**
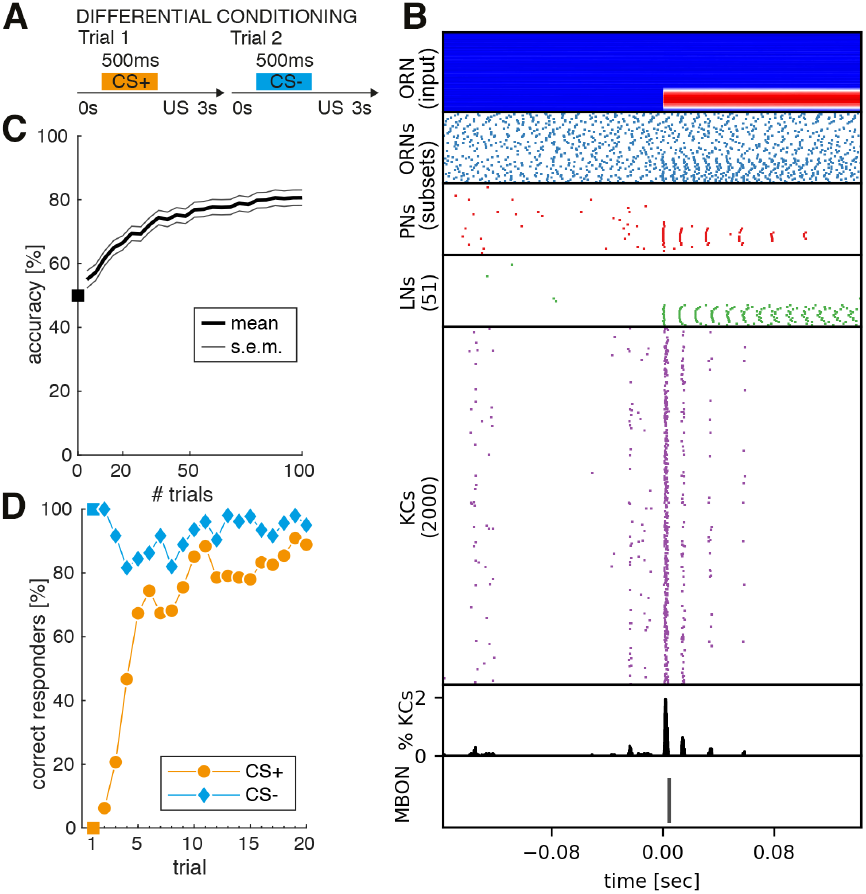
Rapid associative learning with binary behavior response during a classical conditioning task. A: Sketch of a classical conditioning (differential conditioning) paradigm. Two different sensory cues are presented, where one is paired with a reward (CS+), e.g. food, and the other is paired with punishment (CS-). This process is repeated for many trials. After successful learning, the subject is supposed to show a positive binary behavioral response to CS+ cues and no behavioral response to CS-cues. **B:** System response of the olfactory network model (ORN,AL,MB,MBON) in response to a single CS+ stimulus presented during a classical conditioning paradigm. From top to bottom: Input to the model is provided as noisy current injection into the ORN population. Stimulus onset at *t* = 0sec is clearly visible by magnitude increase of injected current for the sub-population of ORNs that belong to the receptor type sensitive to the presented CS+ odor. Stimulus onset is clearly visible by an increase in spiking activity across all populations of the network. For ORNs and PNs only a relevant subset of 60 and 35 neurons of the total population is shown. ORNs (blue raster plot) sensitive to the presented stimulus show an increase in spiking activity over their spontaneous background activity at stimulus onset which slowly attenuates due to the cellular mechanism of spike frequency adaptation (SFA). A similar attenuating effect is visible in the subset of PNs (red raster plot) that belong to the glomeruli of activated ORNs. This is a transitive effect of SFA implemented in the ORNs, as the PNs do not have SFA implemented. The same can be seen in the LN population (green raster plot). The population of 2000 KCs (magenta raster plot) show temporal and spatial sparse response where during spontaneous background activity (Ito et al., 2008) < 1% of all KCs are active and during stimulus onset only about 2% of all KCs are active (cf. histogram in black below magenta KC activity). The transformation of dense code at the ORN population into a temporal and population sparse code at the KC population have been identified as the two main computational principles of the olfactory system. The last row shows response of the plastic mushroom body output neuron (MBON) after being trained in a classical conditioning paradigm. The MBON correctly responds with a single action potential shortly after CS+ cue onset at t = 0sec. **C:** Learning performance of the readout neuron when pairing a single odor cue with a reward (food). The readout neuron is trained to generate a single action potential in response to odor cues associated with CS+ and remain silent for odor cues associated with CS-. A learning step is performed after each trial if and only if the response of the readout neuron was wrong. The median learning dynamics (red) over *N* = 100 independent models show a steep slope up to 80% accuracy within 50 training trials (approximately 25 CS+ and 25 CS-trials). Beyond 50 trials the performance starts to saturate. **D:** Learning dynamics of a binary behavioral response evoked due to presentation of cues. When the readout neuron fires one or more spikes, it is considered to be a positive response and zero spikes are considered a negative (or no) response. A positive response to a CS+ trial or a negative response to a CS-trial is considered to be a correct response. By default the model does not generate any output spike, consequently 100% of the models correctly respond to CS-trials from the beginning (blue). Behavioral response dynamics to CS+ show a steep slope within 10 trials where 80% of the models learn to respond correctly. This result closely matches the few-shot learning dynamics observed in insects, e.g. honeybees, on a similar classical conditioning task.

Population sparse stimulus encoding at the level of KCs is supported by two major factors. First, the sparse divergent-convergent connectivity between the PNs and the 20 times larger population of KCs is the anatomical basis for sparse odor representation (Huerta et al., 2004; Jortner et al., 2007; Litwin-Kumar et al., 2017; Caron et al., 2013; Betkiewicz et al., 2020). Second, lateral inhibition mediated by the LNs in the AL (Wilson, 2013) facilitates decorrelation of odor representations (Wilson and Laurent, 2005; Schmuker et al., 2011; Wilson, 2013; Campbell et al., 2013) and contributes to population sparseness (Luo et al., 2010; Betkiewicz et al., 2020). The sparse code in the KC population has been shown to reduce the overlap between different odor representations (Luo et al., 2010; Lin et al., 2014; Inada et al., 2017) and consequently population sparseness is an important property of olfactory learning and plasticity models in insects (Huerta et al., 2004; Huerta and Nowotny, 2009; Wessnitzer et al., 2012; Ardin et al., 2016; Peng and Chittka, 2017; Müller et al., 2018).

The system response to a single odor presentation in Fig. 2B) demonstrates the transformation of a dense olfactory code at the ORN and PN layers into a population sparse representation at the KC layer where less than < 2% of the total KC population is active at any time during stimulus presentation. This is in good agreement with quantitative estimates in the fruit fly (Turner et al., 2008; Honegger et al., 2011).

### Few-shot learning forms an associative memory of single cues with rewards

Many insects exhibit a rapid learning dynamics when trained in classical olfactory conditioning tasks. They typically acquire high retention scores (test accuracy > 60%) for a binary conditioned response (CR) behavior within only very few trials (e.g. Bitterman et al. (1983); Szyszka et al. (2011); Scheunemann et al. (2013); Pamir et al. (2014)).

Here, we use a classical conditioning paradigm to form associative memories and generate binary CR behavior by training the single MBON in our network (see Fig. 1). We mimic standard experimental lab protocols for classical conditioning (see Fig. 2A). Two different odors are presented as single odor pulses of 500ms duration (see Methods). Multiple independent trials are presented in pseudo-random order where each trial constitutes a single odor stimulus paired with a reward (CS+) or punishment (CS-) shortly after the stimulus presentation.

The system response of the neural network model to a single CS+ stimulus is shown in Fig. 2B. To obtain a neural representation of the odor valence at the MB output (Strube-Bloss et al., 2011; Aso et al., 2014b; Strube-Bloss et al., 2016), the MBON is trained (Gütig, 2016; Rapp et al., 2020) to elicit exactly one action potential in response to a stimulus that is paired with reward (CS+) and zero action potentials when the CS-stimulus is presented (see Methods).

In a first step we quantify the learning performance by considering the accuracy of correctly generated MBON output spikes in response to CS+ and CS-stimuli after each training trial. The average accuracy over *N* = 100 independently trained model instances is shown in Fig. 2C. Learning dynamics of the neural representation of the stimuli show a very steep slope and indicate that memories are formed rapidly with up to 80% accuracy after presentation of 50 (25 × CS+ and 25× CS-) training trials.

Next, we consider the binary CR behavior depending on whether the MBON generates one or more action potentials in response to a stimulus (positive response) or remains silent (negative response). A conditioned response is correct if the MBON generates a positive response to CS+ cues or a negative response to CS-cues. Results are quantified by a behavioral learning curve (Fig. 2 D) representing the median percentage of correctly responding individuals from N = 100 independently trained models.

In the untrained model, and due to randomly drawn initial synaptic weights, the MBON does not generate any action potentials. Thus all models correctly generate negative behavioral responses to CS-trials from the very beginning and consequently zero correct conditioned responses to CS+ stimuli. After presentation of 3 — 5 trials up to 70% of the trained models are able to generate the correct, appetitive CR to the CS+ stimuli. The learning curve saturates after 10 training trials fluctuating around an asymptotic value of 80% correct CRs. This reproduces the rapid learning dynamics of insects in classical conditioning experiments and fits qualitatively and quantitatively to the conditioned response behaviors in honeybees (e.g. Bitterman et al. (1983); Pamir et al. (2011); Szyszka et al. (2011); Pamir et al. (2014)).

We conclude that our statically configured sensory network model with a single plastic readout neuron is capable to successfully form associative memories by few-shot learning, replicating the classical conditioning experiments in the typical lab situation. The computational mechanism of population sparseness implemented in our model increases discriminability of the two different stimuli supporting a rapid learning dynamics and a high accuracy of memory recall.

### Robust dynamic memory recall and background segmentation in complex sensory scenes

Even though sensory systems are highly specialized sub-systems, in general, each sensory system is in charge of solving several sub-problems. This is particularly true for the olfactory system, which is in charge of odor discrimination (Lin et al., 2014; Riffell et al., 2014), segmentation (Grabska-Barwińska et al., 2016) and possibly other perceptual tasks. Thus, whichever neural representation of external stimuli the system uses, this representation should be general to successfully satisfy all of its intended purposes. We now assess our previously trained model, by associative learning of single odor cues in a classical conditioning paradigm, in a new task of recognizing previously associated odors in complex and dynamic olfactory scenes where several different odor cues are present as sketched in Fig. 3 A. We ask the question how robust and general the learned neural representation is, when many cues of different odor sources appear in complex sequences. If successful, we can conclude that our model is able to use a form of compositionality by means of dynamical memory recall of simpler concepts to solve more complex tasks without explicit re-training. The ability of using previously learned concepts to solve novel tasks, in our case without re-training, is also known as *transfer-learning* in the scientific field of machine learning. We draw sequences with an average of five non-overlapping cues, each of random duration between 100ms and 500ms (see Methods). All cues are randomly positioned within *T* = 10s by drawing stimulus onset times from a uniform distribution. Each cue gets randomly assigned to 1 out of up to 5 possible odors. The use of non-overlapping cues follows the rationale presented in Szyszka et al. (2012) who could show that, in nature, filaments originating from different odors do not mix perfectly. The number of possible different odors varies across the 5 different data sets. The data sets are constructed such that the difficulty of odor discrimination increases due to increasing overlap in the ORN receptor activation patterns between odors (similar vs. distinct) and due to an increasing total number of possible odors. Each data-set comprises 200 generated sequences where Fig. 3B shows the system’s response to one example sequence with random occurrences of 3 CS+ and 4 distractor cues. The ORN receptor activation profile of different odors for all 5 odors is shown in fig. 4B.

**Figure 3:**
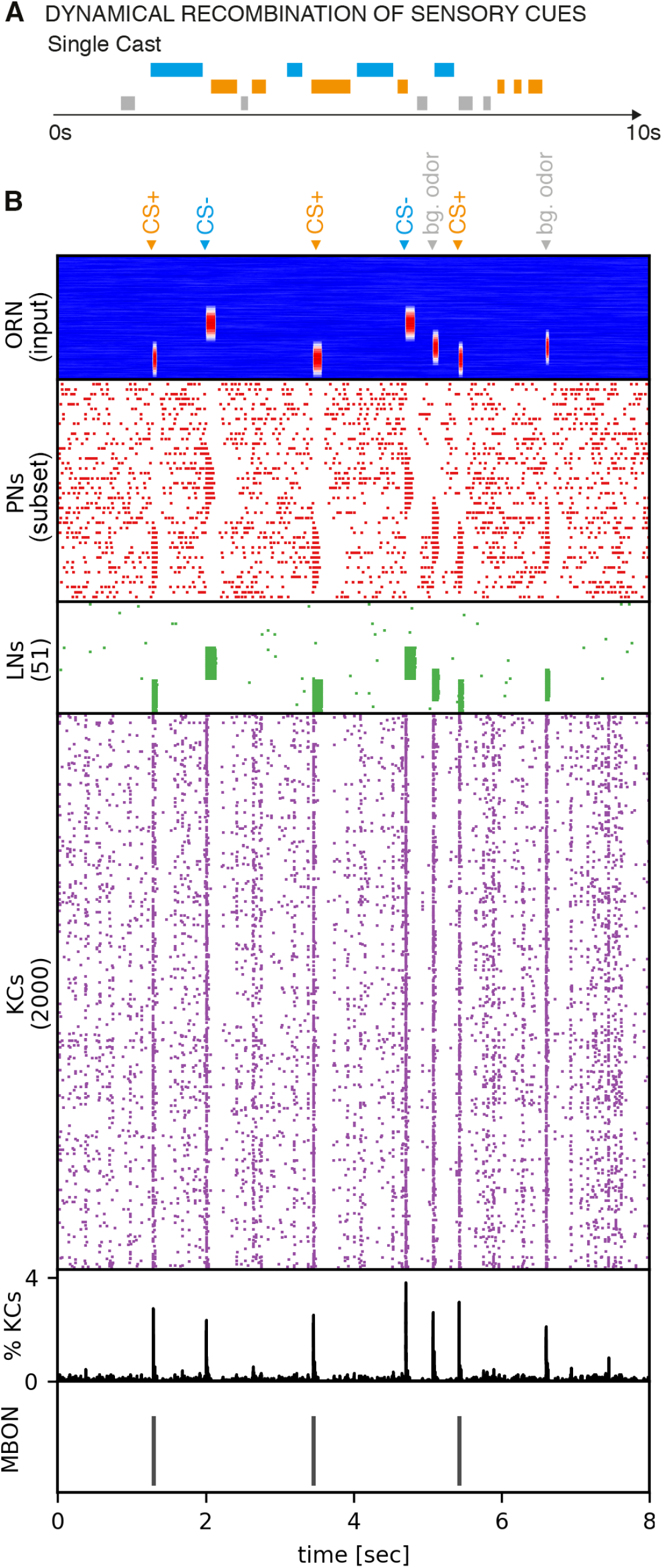
**A:** Sketch of the problem of dynamic memory recall when experiencing multiple cues over time. In a single trial, multiple cues of different odors are experienced over time. The total number of cues over time varies strongly as well as the number of different odors present (from 2 up to 5 in the data-sets used here). Such a sensory experience can be related to natural conditions, where a flying insect encounters many odor filaments over time, e.g. during “cast” behavior employed for foraging. **B** System response of the olfactory network model (ORN input, PN, LN, KC, MBON) in response to a dynamical sequence of cues from 3 different odors. Top to bottom: stimulus profiles of model input as current injection into ORNs; different odors activate PNs that belong to different glomeruli, clearly showing the transitive effect of spike-frequency adaptation implemented in ORNs; activity of LNs evoked by different types of odors; response of KCs with temporal and population sparseness (~ 3% of KCs active during stimulus onset) and spontaneous background activity (Ito et al., 2008); response of the readout MBON with one action potential to each CS+ related cue whereas no action potentials are generated for CS-cues and all other distractor cues.

**Figure 4:**
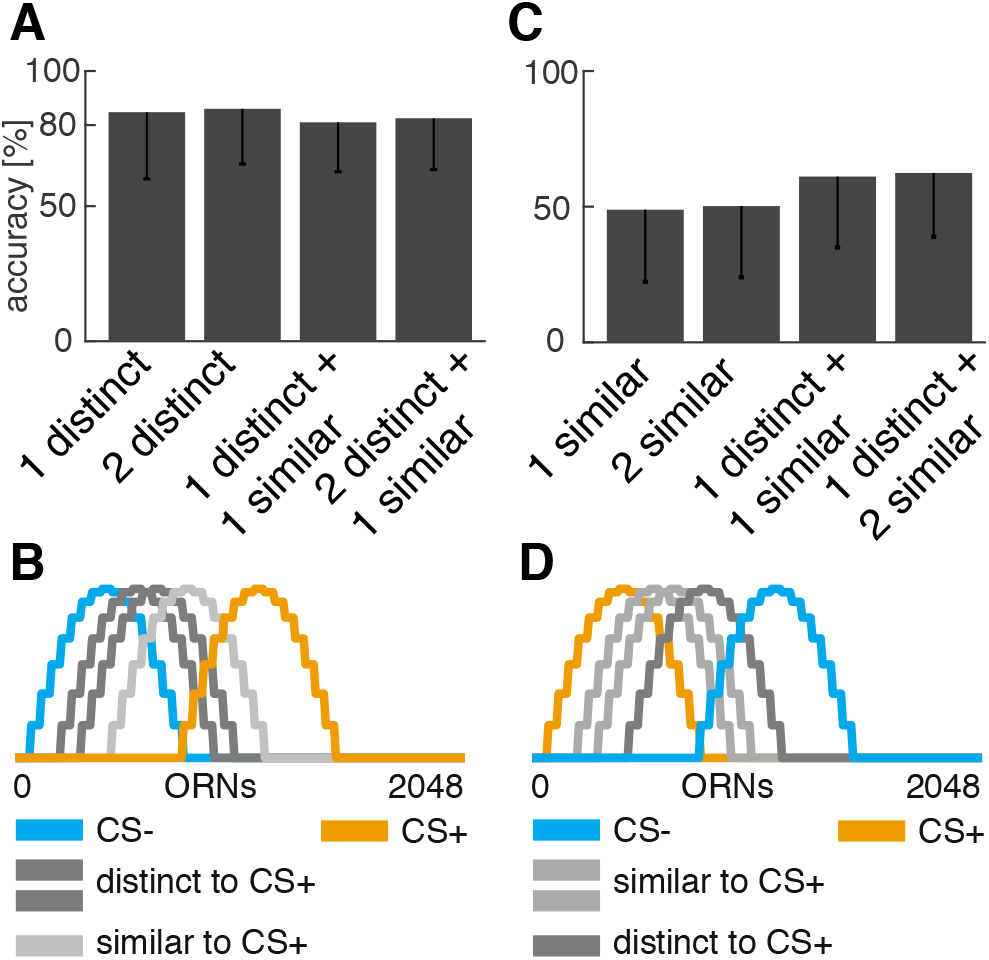
Fast and accurate dynamic memory recall of multiple cues using previously learned associations. **A:** Results when testing the trained model on the problem of dynamic memory recall of multiple cues. Each of the five different data sets is quantified by accuracy of correctly estimated trials. A single trial constitutes a sequence of multiple sensory cues and is considered correct when the number of CS+ cues within that sequence are correctly predicted by a single action potential per each CS+ cue. Cue distributions are modeled as Poisson-like events. The difficulty increases with each data set in terms of the number of different odors present in a sequence and similarity (ORN stimulus response overlap) of pleasant vs. unpleasant vs. background odors. The results show that the model can successfully solve this new and more complex problem without additional training. **B:** Similarity of odors in terms of overlap in ORN stimulus response profiles for the 5 different odors. Here, the odor associated with CS+ is distinct from the odor associated with CS-(no overlap). Out of the 3 distractor odors each has a slightly larger overlap with the CS-odor and is considered to be more similar to CS-. **C+D:** The same type of experiment, but with different ORN response profiles. Now the CS+ odor is very similar to all background odors and renders the problem more difficult. All models have been trained in the same classical conditioning task but using the ORN response profiles shown in panel B,D. Afterwards they are tested on the five generated data sets that contain recombination of multiple cues without retraining. Results show, that the model generalizes reasonably to this problem despite the strong similarity of CS+ and distractor odors.

The objective in this task is a neuronal response with a single action potential to each cue occurrence that belongs to the positively associated odor (CS+). For all other cues, CS-as well as any distractor cue, no action potential should be generated. The overall response to a sequence is considered to be correct if and only if the number of action potentials generated by the readout neuron is equal to the number of CS+ cues. Results are shown in fig. 4A. We find that our previously trained model can successfully generalize to this new task and reaches ~80% accuracy across all data sets, independent of their complexity. It is thus able to use the previously learned neural representation of single cues and dynamically recalls memories to respond correctly to complex sequences of stimuli of different length. We further find, that our model solves the problem of odor vs. background segmentation (Grabska-Barwińska et al., 2016), by reliably responding to CS+ cues only as quantified by the accuracy of correctly recalled CS+ cues in the sequences.

In a second step we performed the same task, but changed the odor associated with CS+ to have large overlap with all 4 distractor odors in terms of ORN receptor activation profile (Fig. 4D). We used the same conditioning paradigm with single cues to train the model and test it on the generated sequential data sets (Fig. 4C). We find that, despite the large overlap of the receptor profiles of CS+ odor and the distractor odors, our model is still able to generalize reasonably well with accuracy of 50% correctly detected sequences across all data sets.

We conclude that our model can use a form of compositionality to successfully implement transfer learning without re-training by robustly generalizing memory recall from single cues to complex sequences of multiple cues and distractor cues of variable cue duration. Our results hold for arbitrary distributions of cues in time. As a by-product the model solves the problem of odor background segmentation. The results presented in 4A+D suggest, that the functional role of temporal sparseness is to generate precise temporal memory recall up to very short time scales (i.e. for short cue durations and high frequency of cue occurrences).

### Sensory evidence accumulation informs motor control for foraging

We now consider the situation of foraging within a natural environment (fig. 5A). The objective is to locate the food source which emits an attractive odor (CS+) by utilizing the sensory cues present in its turbulent odor plume. We show that cast & surge behavior can emerge by accumulation and exploitation of sensory evidence of sequentially experienced individual cues.

**Figure 5:**
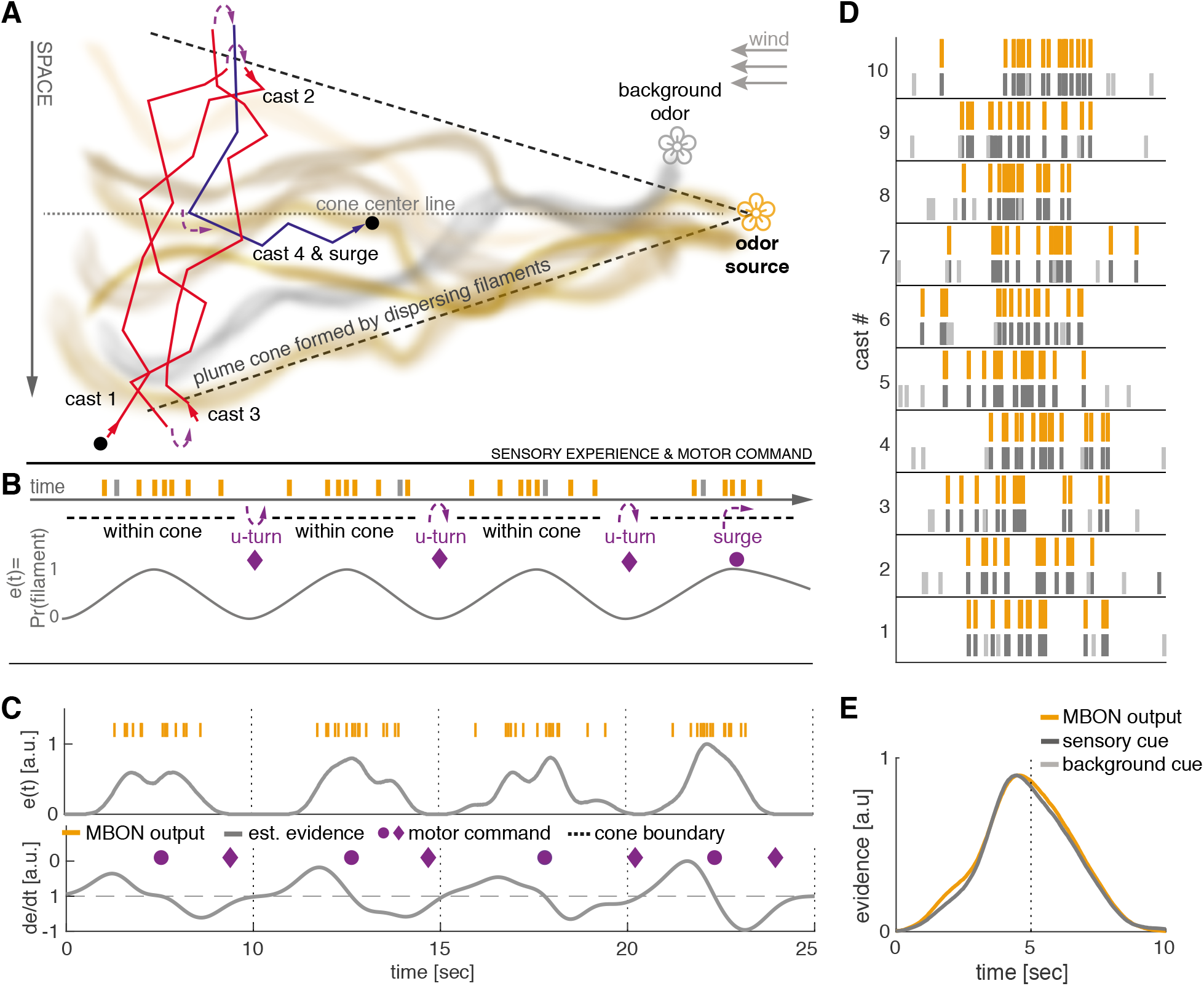
Dynamical sensory processing and motor control serving chemotaxis A: Sketch of a typical olfactory experiment setup in a wind tunnel with a pleasant odor source (orange flower) and a second distractor source (gray flower). Due to turbulence, the odor molecules emitted by a source form dispersing, intermittent filaments within a cone-like boundary that constitutes the odor plume. The plume is modeled as Gaussian distributed filaments. A behaving model insect (here *Drosophila melanogaster*) performs stereotypic “cast&surge” behavior to locate the food source. This constitutes alternating between scanning crosswind and U-turning after running past the plume cone boundary where no filaments are present. Eventually after several casts (here 3) it surges upwind until it loses track of the plume cone and starts over. **B:** Filament encounters during this behavior result in brief on/off, sequential stimuli of the olfactory system. The probability of encountering filaments is > 0 within the plume and zero outside of the plume. Sensory evidence *e*(*t*) can be viewed as a likelihood function of filament encounters, that increases towards the plume’s center line and is zero outside of the plume. The properties of this function can be used to generate optimal motor commands for chemotaxis. **C:** Evidence *e*(*t*) and derivative 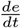 over 4 simulated casting trajectories estimated by low-pass filtering the MBON spiking activity. <u>U-turn</u> motor commands are generated when *e*(*t*) runs below a fixed threshold (0.01) and <u>surge</u> motor commands are generated when 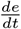 turns negative. The motor commands generated by this model closely match the optimal commands as sketched in panel B. **D:** Spiking activity of the MBON (orange) in response to 10 casting trajectories. The MBON reliably predicts the true sensory cues of simulated filaments (dark gray) and ignores background cues (light gray). **E:** Smooth PSTH computed over 10 casting trials recovers an accurate estimate of the true underlying sensory cue distribution simulated from a gaussian distribution.

For this task, we assume that the occurrence of filaments within a cross-wind slice of the plume are approximately Gaussian distributed. This is a reasonable assumption, particularly in a wind-tunnel setting with laminar flows, as typically used in experimental settings (van Breugel and Dickinson, 2014; Sehdev et al., 2019). However, our model is general and works independent of the actual distribution of filaments. When the insect performs a cast through the plume, it encounters filaments as short-lived discrete, sequential events where each encounter represents a single sensory cue (see sketch in Fig. 5B). Thus, to simulate the sensory experience during casting we generate sequences of cues and distractors, where cue onsets for CS+ odor are drawn from a Gaussian distribution and onsets of distractor cues from a uniform distribution (see Methods). We further assume that the subject has already formed an association of food with the attractive odor, either through learning (e.g. classical conditioning, cf. Fig.2) or through some genetically predetermined innate valence. To this end we again use the trained model from the classical conditioning task above without any further re-training.

We simulate 4 consecutive casting trajectories where the agent senses odor cues of sequentially experienced filament encounters. Ongoing accumulation of sensory evidence (Fig. 5C) by low-pass filtering of the readout neuron’s output assumes positive values shortly after entering the plume cone and further increases while approaching the plume’s center line. When travelling beyond the center line sensory evidence slowly decreases until the agent leaves the plume cone boundary. When sensory evidence drops to zero and after a fix delay, the agent initiates a U-turn motor command to perform another casting trajectory. Responses from our model’s readout neuron precisely follow the ground truth of CS+ odor cues as shown by 10 random casting trajectories in Fig. 5D. Performing analysis by averaging of sensory evidence across these 10 casting trajectories yields an average evidence (Fig. 5E) that resembles the underlying, true Gaussian profile of the simulated filaments. We thus find that the average estimate produced by our model closely matches the ground truth distribution.

We conclude that the model output provides an accurate and robust estimate of sensory evidence that can be used to reason about a plume’s spatial extend and center line. Both information are crucial to generate appropriate motor commands for U-turn and upwind surge behavior, necessary to successfully execute a cast & surge strategy. Apart from the existence of filaments inside a plume and absence outside a plume’s cone, our model does not make any specific assumption regarding the plume’s structure and statistics. It thus provides a generic mechanism implemented in a neural system to perform cast & surge behavior during foraging flights.

## 3. Discussion

### Distinct functional roles for population and temporal sparse stimulus encoding

Population sparseness improves discriminability of different stimuli to facilitate associative learning. This has been demonstrated in theory and experiment (Lin et al., 2014; Litwin-Kumar et al., 2017; Caron et al., 2013; Assisi et al., 2020; Nowotny et al., 2003; Jortner et al., 2007). We have shown, that our neural network model implements this feature in a biologically realistic way and our results confirm the functional role of population sparseness to support rapid and robust memory acquisition through associative learning. Experimental Liu and Davis (2008); Lin et al. (2014) and theoretical (Bennett et al., 2019; Assisi et al., 2020; Litwin-Kumar et al., 2017) studies in the fruit fly strongly suggests that inhibitory feedback through the anterior paired lateral (APL) neuron improves population sparseness in the KC population. The APL is a GABAergic neuron that broadly innervates the KC population and likely receives input from KCs in the MB output region. Inhibitory feedback from MB output onto MB input has also been demonstrated in other species (Rybak and Menzel, 1993; Papadopoulou et al., 2011) and blocking of feedback inhibition in the MB reduced population sparseness in the honeybee (Froese et al., 2014). Including an inhibitory feedback loop in our model would further improve robustness of population sparseness and thus not change the our core findings.

Our model demonstrates how temporal sparseness can be exploited to generate short patterned signaling of cue identity. This enables perception of high temporal stimulus dynamics. In our model this is achieved independently of the duration of individual stimulus incidents and their distribution in time and makes temporally precise and robust sensory evidence available. It allows for the ongoing computation of derived estimates such as cue distributions or changes in cue density. Maintaining temporally sparse representations mechanistically supports the principle of compositionality (or Frege principle (Hintikka, 1984)), where an atomic stimulus entity is represented and can be learned by the readout neuron before processing this output. For example by estimation of densities or recombination with other entities to form composite perception or memory read-out. The temporal stimulus dynamics remains intact throughout the system even after learning of stimulus relevance. Thus valence is encoded with the same dynamics and faithfully captures occurrences of relevant cues. This allows compression of code to relevant stimuli while retaining full stimulus dynamics of the external world. Compression of code along the sensory processing pipeline is particularly relevant for small-brained animals like insects, which need to economize on their neuronal resources.

### Odor-background segregation: a joint effect of temporal and population sparse cue representation

The task presented in Fig. 3 implicitly addresses the issue of odor-background segregation. This refers to the problem that in nature cues of multiple odors of different sources are present, either in terms of mixtures or stimulus onset asynchrony due to turbulent conditions (Szyszka et al., 2012; Sehdev and Szyszka, 2019; Grabska-Barwińska et al., 2016; Chakroborty et al., 2016; Deisig et al., 2010; Capurro et al., 2012). For behavior it is relevant to reliably isolate and detect the relevant cues from any background or distractor cues. The results presented in Fig. 4 show that this works nicely in our system. This is achieved by exploiting the joint effect of temporal and population sparseness. Optimal discrimination of cue representation is guarantied by population sparseness and temporal precision by means of temporal sparseness. Our plastic output neuron requires population sparseness for learning and the plasticity rule (Gütig, 2016; Rapp et al., 2020) allows for temporally precise memory recall. We predict that our model can solve the challenge of odor-background segregation.

### Rapid learning within few trials

The ability of insects to quickly form associative memories after 3-5 trials has been demonstrated experimentally (Szyszka et al., 2011; Pamir et al., 2014). In Nowotny et al. (2005) a model has been shown to perform single-shot learning to discriminate odors. However, in general few-shot learning remains a difficult task for computational models (Delahunt and Kutz, 2019; Jiang and Litwin-Kumar, 2019). We find that, when compared with learning dynamics data of real insects (Szyszka et al., 2011; Pamir et al., 2014) our model is able to show realistic learning dynamics that matches with the experimental observations. Due to frequent changes in the environment it might be a better strategy to tradeoff fast and reasonable accurate learning against slow and high precision learning. Additionally, acquisition of training samples might be costly or they generally occur very sparsely.

Few-shot learning capabilities are also an active area of research in machine learning. Particularly current deep learning methods require massive amounts of training samples to successfully learn a classification model. For example the popular benchmark data sets, ImageNet (Deng et al., 2009) and CIFAR10 (Krizhevsky, 2009), for image classification contain 14 million and 60 million sample images. The *Google News dataset* used to train language models contains 100 billion words and learning to play the Space Invaders Atari game by deep reinforcement learning requires sampling of > 500.000 game frames from the environment. Clearly, these are numbers a biological organism cannot afford to accumulate. In fact, all animals have been shown to be able to perform few-shot learning which likely is a fundamental skill for survival. We have demonstrated that our neurobiologically motivated approach using spike-based computations is capable to perform few-shot learning with similar speed as real insects to establish an odor-with-reward association. We further showed, that our model can perform transfer learning to novel, complex combinations that have not been part of the training data.

### Innate vs. learned behavior

Cast & surge behavior belongs to the innate behavioral repertoire of air-borne insects and emerges from a set of sensorimotor reflexes (van Breugel and Dickinson, 2014). It can be considered as a base strategy wich guarantees survival. The base system can be modulated and improved throughout an animals lifespan by experience-based learning. On the other hand, strategies that emerge solely from learned behavior might require re-training whenever the environment changes. This can be costly in terms of energy, acquisition of samples to learn from or a short life span in general. Here, we assume, that our readout neuron is tuned to a pleasant odor. In the present work this tuning is learned (adaptive process) as we have shown with the classical conditioning task. However, the tuning can generally be learned by other mechanisms, e.g. reinforcement learning. We demonstrated that the existence of such a tuned neuron allows cast & surge foraging behavior to emerge.

There are other ways how such a tuned neuron can come about, for example due to genetically predetermined wiring or during development from larval to adult stage. The cast & surge behavior can be executed on innately valenced olfactory cues and our suggested model for motor control during cast & surge (Fig. 5A+B) also works for innate valenced stimuli. Learning is important to adapt behavior to changing environmental circumstances and associative learning provides a means to learn new valences on demand in such situations. Our model learns odor valences at the mushroom body output and it has been shown that MBONs signal odor valence (Hige et al., 2015; Owald et al., 2015; Aso et al., 2014b; Perisse et al., 2016; Strube-Bloss et al., 2016, 2011). We suggest that this valence is then used downstream to execute higher level functions of motor control. At this processing stage it might be integrated with innate valences and other necessary sensory input (Fry et al., 2009; Ache et al., 2019; van Breugel and Dickinson, 2014; Saxena et al., 2018) to form behavioral decisions.

### Implications for other sensory systems

Sparse stimulus encoding has been identified as a powerful principle used by higher order brain areas to encode and represent features of the sensory environment in invertebrate (Perez-Orive et al., 2002; Szyszka et al., 2005; Turner et al., 2008; Ito et al., 2008; Honegger et al., 2011) and vertebrate (Hromádka et al., 2008; Vinje, 2000; Wolfe et al., 2010; Isaacson, 2010) systems. Sensory systems with similar coding principles may share similar mechanisms when it comes to learning and multi-modal sensory integration. The mushroom body is a center for integration of multi-modal sensory information. Thus our model can be extended to incorporate multi-modal input from other sensory systems. It is known that olfactory search and foraging strategies do not solely rely on olfactory cues, but require additional sensory information from at least visual cues (Fry et al., 2009; Ache et al., 2019; van Breugel and Dickinson, 2014; Saxena et al., 2018) and wind direction (Wolf and Wehner, 2000; Bhandawat et al., 2010; Álvarez-Salvado et al., 2018; Suver et al., 2019). Extending our model to include additional sensory processing systems for vision and wind direction can provide a comprehensive functional model to study foraging, olfactory search and navigation.

### Potential improvement through multiple readout neurons

Our current approach only comprises the simplest case of a single readout neuron. This model can be extended to multiple readout neurons. Different readout neurons can be tuned to different odors or groups of odorants. This would allow foraging for different types of food sources and further be useful for multi-modal sensory integration and learning of valences of multiple odors. Another way to use multiple readout neurons is to create an ensemble learning model. Particularly, one can perform bootstrap aggregation (bagging) (Breiman, 1996) to decrease variance of predictions. With this technique, multiple, independent readout neurons are trained for the same target and their outputs are averaged to produce a single output. This approach can be useful when the level of noise increases due to different input models used to drive the network. Another possible extension is to use a single readout neuron to code for multiple odors by associating different number of action potentials to different odors (e.g. 2 or 3). The choice of model for the readout neuron and the plasticity rule allows to do this (Gütig, 2016).

### Top-down motor control and lateral horn

In the current work, generation of motor commands is not computed within the neural network. Integration of sensory evidence is modeled by low-pass filtering of the readout neuron’s spike train and its derivative is numerically estimated. In Lundstrom et al. (2008) it has been shown, that a single compartment Hodgkin-Huxley neuron can operate in two computational regimes. One is more sensitive to input variance and acts like a differentiator while in the other regime it acts like an integrator. Similarly Ratté et al. (2015) has shown that the subthreshold current of neurons can encode the integral or derivative of their inputs based on their tuning properties. This could serve as basis for estimating the low-pass filtered sensory evidence and its derivative using neural computations. The turning behavior could be implemented by an always-on neuron serving as a central pattern generator for the motor signal. This neuron can be inhibited by the activity of the readout neuron and only becomes active when no sensory cues are present anymore, which happens shortly after leaving the plume cone. Another option could be to use the cellular mechanism of spike-frequency adaptation to initiate a fast turning movement which slowly decays with a fixed time constant.

### Relevance for machine learning and artificial intelligence

Learning and building artificial, intelligent agents capable of interacting with their environment are major objectives in the field of machine learning (ML) and artificial intelligence (AI). Deep artificial neural networks (LeCun et al., 2015; Schmidhuber, 2015) have demonstrated great success over the recent years. Particularly, in the domains of image recognition (Krizhevsky et al., 2012; Simonyan and Zisserman, 2014), natural language processing (Pennington et al., 2014; Bahdanau et al., 2014; Mikolov et al., 2013; Vaswani et al., 2017; Devlin et al., 2018) and deep reinforcement learning (Mnih et al., 2015; Lillicrap et al., 2015). Despite their success, when applied to agent-based systems, their major drawback becomes evident. They are very specific, single-purpose perceptual systems and poorly generalize to new tasks or changes in an agent’s environment (non-stationarities). A few methods to overcome this problem have been proposed, this includes re-training on new tasks and transfer-learning. In the context of deep learning this refers to the method of training a base network on features that are general to all tasks. Afterwards the pre-trained base network is used and the learned features are repurposed to only train a classification layer on the new tasks. However, it turned out that re-training brings up another weakness of deep neural networks, catastrophic forgetting (Kirkpatrick et al., 2017). This term refers to the fact, that after a model has been trained on one task and gets re-trained on a 2nd task, it will completely forget everything it has learned on the previous task. In this work we used a method similarly to the latter approach of transfer-learning but without any additional retraining and we used spike-based learning based on an improved implementation (Rapp et al., 2020) of the Multispike Tempotron (Gütig, 2016). We predict that spike-based methods inspired by biological learning will become increasingly important for artificial intelligence.

## Acknowledgments

This research is supported by the German Research Foundation (grant no. 403329959 to MN) within the Research Unit ‘Structure, Plasticity and Behavioral Function of the Drosophila mushroom body’ (DFG-FOR 2705, www.uni-goettingen.de/en/601524.html).

## Author Contributions

Conceptualization, H.R., MP.N.; Methodology, H.R and MP.N.; Writing – Original Draft, H.R., Writing – Review and Editing, H.R. and MP.N.

## 5. Methods

### LEAD CONTACT AND MATERIALS AVAILABILITY

This study did not generate new unique reagents.

### METHOD DETAILS

#### Spiking network model

All neurons of the olfactory network are modeled as conductance-based leaky integrate-and-fire neurons with spike frequency adaptation (SFA). Specifically, the membrane potential follows the dynamical current balance equation 1. On threshold crossing a hard reset of the membrane potential is performed by 2. SFA is modeled as outward current by term 4 of equation 1. Strength of the adaptation current is modeled by a constant (b) decrease on each threshold crossing. Input to the model is modeled as direct, time-dependent current injection of shot-noise to all ORN neurons by the term *I_stim_*(*t*). All simulations of the network are carried out using BRIAN2 (Stimberg et al., 2019) simulator. The membrane potential of each neuron within a population is initialized randomly ∈ [*V_rest_, V_threshold_*]. To avoid any artifacts the network is brought to equilibrium by driving the network for 2 sec with background activity only before starting the actual simulation.

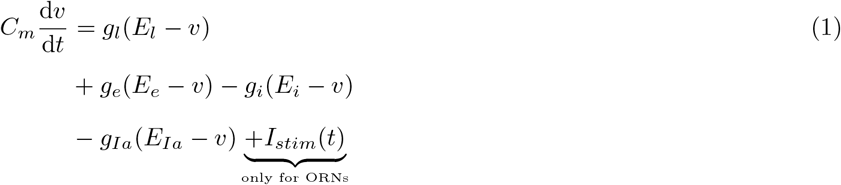

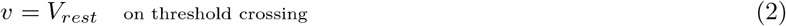

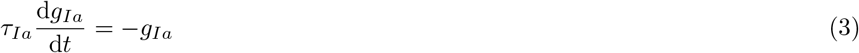

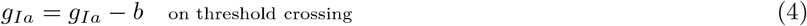

For this work the number of neurons within each layer and connectivity schemes are chosen to match the numbers found in the adult *Drosophila melanogaster* (Takemura et al., 2017; Aso et al., 2014a). Our model comprises 2048 explicitly modeled olfactory recepter neurons (ORNs) organized in 52 different receptor types. ORNs of the same receptor type converge onto the same Glomerulus (52) by feedforward excitatory synapses. Each Glomerulus is formed by a projection neuron (PN) and local interneuron (LN). LNs provide lateral inhibition to all other PNs and LNs. PNs randomly project to a large population (2000) of Kenyon cells (KC) with excitatory synapses such that each KC on average receives input from 6 random PNs. This sparse random convergence implements population sparse responses. The single, plastic mushroom body output neuron is fully connected to all KCs.

We used the cellular mechanism of spike-frequency adaptation (SFA) to achieve temporal sparseness. ORNs are configured to have slow and weak spike-frequency adaptation in accordance with experimental findings (Nagel and Wilson, 2011; Bhandawat et al., 2007; Krofczik et al., 2009). For PNs and LNs SFA has been turned off and KCs are set to produce fast and strong adaptation currents (Wustenberg et al., 2004; Demmer and Kloppenburg, 2009). The property of temporal sparseness can also be achieved by an alternative implementation through feedback inhibition as proposed by Assisi et al. (2020) and Bennett et al. (2019).

The synaptic weights of all connections within the network have been manually determined such that an average background firing rate of 8 – 10 Hz is achieved in the LN population. All parameter values used for neurons of each population are summarized in tables 1,2,3,4. Other parameters used to setup the network (time constants, synaptic weights) are summarized in table 5.

**Table 1:** Neuron model parameters used for ORN population.

| ORN |  |  |
| --- | --- | --- |
| Parameter | Value | Unit |
| V_threshold | -57.00000 | mV |
| b | 2.00000 | nS |
| tau_Ia | 1.00000 | s |
| V_rest | -70.00000 | mV |
| gL | 28.95000 | nS |
| EL | -70.00000 | mV |
| C | 289.50000 | pF |

**Table 2:** Neuron model parameters used for PN population.

| PN |  |  |
| --- | --- | --- |
| Parameter | Value | Unit |
| C | 289.50000 | pF |
| tau_Ia | 1.00000 | s |
| V_rest | -70.00000 | mV |
| b | 0.00000 | S |
| gL | 28.95000 | nS |
| EL | -70.00000 | mV |
| v_threshold | -57.00000 | mV |

**Table 3:** Neuron model parameters used for LN population.

| LN |  |  |
| --- | --- | --- |
| Parameter | Value | Unit |
| C | 289.50000 | pF |
| tau_Ia | 1.00000 | s |
| V_rest | -70.00000 | mV |
| b | 0.00000 | S |
| gL | 28.95000 | nS |
| EL | -70.00000 | mV |
| V_threshold | -57.00000 | mV |

**Table 4:** Neuron model parameters used for KC population.

| KC |  |  |
| --- | --- | --- |
| Parameter | Value | Unit |
| C | 289.50000 | pF |
| tau_Ia | 50.00000 | ms |
| V_rest | -70.00000 | mV |
| b | 5.00000 | nS |
| gL | 28.95000 | nS |
| EL | -70.00000 | mV |
| V_threshold | -57.00000 | mV |

**Table 5:** Other parameters used to configure the model (time constants, synaptic weights, etc.).

| Network |  |  |  |
| --- | --- | --- | --- |
| Parameter | Value | Unit | Description |
| tau_syn_i | 10.0 | ms | time constant inhibit. synapses |
| tau_syn_e | 2.0 | ms | time constant excit. synapses |
| tau_ref | 5.0 | ms | time constant refractory period |
| w_orn_pn | 1.1282 | nS | synaptic weight for ORN-PN synapses |
| w_orn_ln | 1.0 | nS | synaptic weight for ORN-LN synapses |
| w_ln_pn | 2.5 | nS | synaptic weight for LN-PN inhibitory synapses |
| w_pn_kc | 14.0 | nS | synaptic weight for PN-KC synapses |

#### Stimulus response profile of ORNs

The stimulus response profile of ORNs is determined by the ORN tuning curves. We follow a similar method as used in Betkiewicz et al. (2020) where cyclical tuning over receptor types is modeled as half period sine waveforms. Our model comprises *N_type_* = 52 receptor types and supports 52 different stimuli (e.g. different odors). Where *k_type_* refers to the receptor type index (∈ [0, 51]) and *k_odor_* to the stimulus index (∈ [0, 51]). *N_orn_* = 15 determines the number of receptor types activated by a stimulus. The tuning strength *r* of the ORNs can be computed as 0.5 cycle of a sine wave with peak amplitude *r_max_* = 1. In the present work all tuning profiles are normalized to have a peak amplitude of 1.

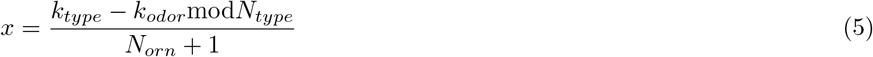

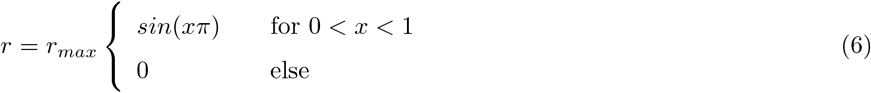

#### Model input

Input to the mushroom body model is modeled as time-dependent, direct current injection into all ORN neurons. In the absence of any stimuli ORNs exhibit spontaneous activity (Nagel and Wilson, 2011). The model input thus consists of spontaneous background activity and stimulus related activity. To generate the background activity, a current time-series is generated for each ORN by simulating shot noise. For each ORN neuron, background activity events are generated from a Poisson process with high rate (λ = 300) (independent Poisson processes are drawn for each individual neuron). Events of the Poisson process are filtered by a low-pass filter with *τ* = 0.6 sec. Using this shot-noise model is consistent with experimental findings of odor transduction at the ORNs (Nagel and Wilson, 2011). To induce stimulus related activity to this time-series of ORN *j* it is multiplied point-wise with a stimulaton protocol time-series *S_j_* (t) which is rescaled by a constant determined by the tuning strength (*r_j_* ∈ [0,1]) to the specific odor of the ORN. This results in a current time series where during stimulus the current magnitude is increased proportional to the ORNs tuning strength and otherwise remains at the magnitude of the background activity.

We define a stimulation protocol function *s*(*t*), which is a step function taking on the value 1 at all time points *t* where a stimulus or sensory cue is active. For each ORN a rescaled instance of the stimulation protocol is defined as *S_j_*(*t*) = *r_j_s*(*t*), where the scaling parameter *r_j_* ∈ [0,1] is given by the stimulus response profile (eq. 6) of the ORN to the specific stimulus.

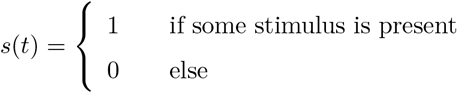

#### Sequences of sensory cues

Each sequence has a duration of 10 seconds. Sequences of sensory cues are generated by drawing the total number of cues within a single sequence from a Poisson distribution with mean λ = 8. Onset times of the cues between 0 and 10 seconds are drawn from a random uniform distribution and it is assured that there is no temporal overlapp between cues. A stimulus relates to a single sensory cue and its duration is drawn uniformly between [1, 200] milliseconds. Finally, each sensory cue is associated with a random odor drawn from a fixed set of possible odors (random sampling with replacement and equal probability). This results in sequences with random number of sensory cues, random onset, random duration and randomized odor and distractor combinations.

#### Model of sensory cues within (gaussian) plume

The same procedure is used as above to simulate the experience of sensory cues during a single casting trajectory within a turbulent odor plume. The number of pleasant cues experienced in a casting trajectory is drawn from a Poisson distribution with mean λ = 14. The cue onset times are drawn from a gaussian distribution with *μ* = 5, *σ* = 1.5. The number of distractor cues is drawn from a Poisson distribution with mean λ = 5 and are distributed uniformly in time. Duration of both, pleasant and distractor cues, is drawn uniformly between [100, 500] ms. In total 200 different casting trajectories have been generated using this procedure.

#### Readout Neuron & Learning rule

To fit the readout neuron to the stimuli such that it generates 1 spike for pleasant odor stimuli (CS+) and 0 spikes for any other stimuli (CS-) we use a modified implementation of the Multispike Tempotron Gütig (2016); Rapp et al. (2020). Thus, the readout neuron is modeled as voltage-based leaky integrate-and-fire neuron with soft reset following the dynamical equation 7. Incoming spikes evoke exponentially decaying post-synaptic potentials. When the membrane potential reaches the spiking threshold at some time *to* an output spike is generated and the membrane potential is reset by the last term of equation 7.

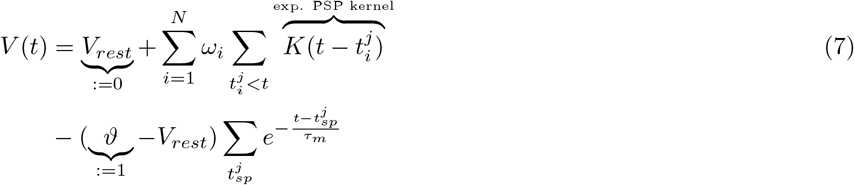

The dynamical equation can be decomposed into two parts, the unreset sub-threshold potential *V*_0_(*t*) (eq. 8) minus the remaining terms for the soft-reset. The neuron is trained to generate 1 spike for pleasant odor stimuli (CS+1) and 0 spikes for any other stimuli (CS-). To fit the desired neural code, a training step is performed after each stimulus presentation. A training step is performed only if the number of spikes generated in response to a stimulus was not correct. The training target is given by the difference between number of output spikes the model generated and the number of output spikes associated with the stimulus. We denote the desired critical threshold value, the voltage value that generates *d* =1 spike, as *ϑ*\* and the time point where this voltage value is reached by *t*\* (more generally: the critical threshold value to generate *d* spikes). We briefly sketch the idea and intuition of the Multispike Tempotron learning rule. For detailed derivation of the rule we refer to Gütig (2016) and the section <u>The *ϑ*\* gradient</u>. The Multispike Tempotron training algorithm works by differentiating the membrane potential of the the critical threshold wrt. to the synaptic weights 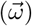. This can be done since *ϑ*\* is a regular voltage value, that can be expressed by the neuron’s dynamical equation (7), with the special identities shown in equation 10. This allows to take the full derivative as shown in equation 11.

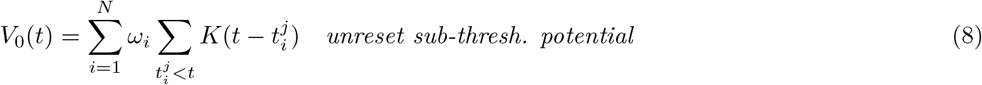

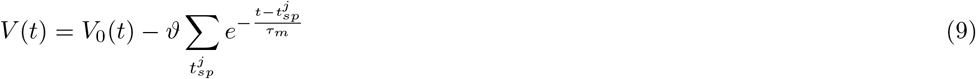

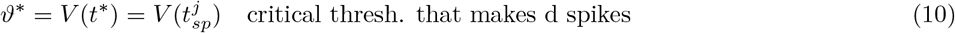

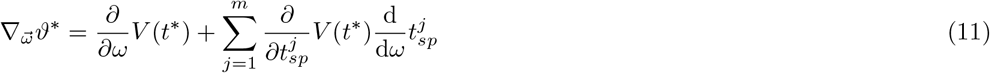

The gradient of the critical threshold with respect to a single synapse *i* is given by equation 12.

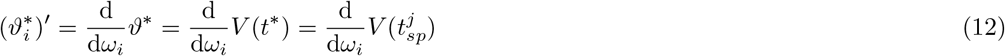

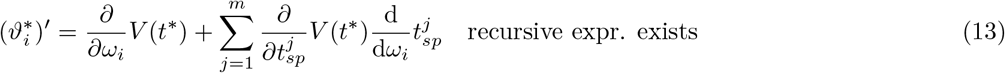

### QUANTIFICATION AND STATISTICAL ANALYSIS

Spiking network simulations have been performed using BRIAN2 and Python 3.6. Model fitting, data analysis and visualization has been done in MATLAB 2018a and partly in Python using matplotlib.

### DATA AND CODEAVAILABILITY

Code and data sets will be made available through our github profile at: https://github.org/nawrotlab

